# Insights into the Absence of Lymphoma Despite Fulminant Epstein-Barr Virus Infection in Patients with XIAP Deficiency

**DOI:** 10.1101/2025.01.17.633616

**Authors:** Yizhe Sun, Janet Chou, Kevin Dong, Steven P. Gygi, Benjamin E. Gewurz

**Affiliations:** Division of Infectious Diseases, Department of Medicine, Brigham and Women’s Hospital, Boston, Massachusetts, United States of America; Harvard Program in Virology, Harvard Medical School, Boston, Massachusetts, United States of America; Division of Immunology, Department of Pediatrics Harvard Medical School, Boston Children’s Hospital, Boston, Massachusetts, USA; Department of Cell Biology, Harvard Medical School, Boston, Massachusetts, United States; Center for Integrated Solutions for Infectious Diseases, Broad Institute of Harvard and MIT, Cambridge, Massachusetts, United States of America; Department of Microbiology, Harvard Medical School, Boston, Massachusetts, United States of America

## Abstract

X-linked Lymphoproliferative Syndromes (XLP), which arise from mutations in the *SH2D1A* or *XIAP* genes, are characterized by the inability to control Epstein-Barr Virus (EBV) infection. While primary EBV infection triggers severe diseases in each, lymphomas occur at high rates with XLP-1 but not with XLP-2. Why XLP-2 patients are apparently protected from EBV-driven lymphomagenesis, in contrast to all other described congenital conditions that result in heightened susceptibility to EBV, remains a key open question. To gain insights, we cross-compared newly EBV infected versus immune stimulated B-cells from XLP-2 patients or upon XIAP CRISPR knockout, relative to healthy controls. XIAP perturbation impeded outgrowth of newly EBV-infected primary human B-cells, though had no impact on proliferation of B-cells stimulated by CD40 ligand and interleukin-21 or upon established EBV-immortalized lymphoblastoid cell lines (LCLs). B-cells from XLP-2 patients or in which XIAP was depleted by CRISPR editing exhibited a markedly lower EBV transformation efficiency than healthy control B-cells. Mechanistically, nascent EBV infection activated p53-mediated apoptosis signaling, whose effects on transforming B-cell death were counteracted by XIAP. In the absence of XIAP, EBV infection triggered high rates of apoptosis, not seen with CD40L/IL-21 stimulation. Moreover, inflammatory cytokines are present at high levels in XLP-2 patient serum with fulminant EBV infection, which heightened apoptosis induction in newly EBV-infected cells. These findings highlight the crucial role of XIAP in supporting early stages of EBV-driven B-cell immortalization and provide insights into the absence of EBV+ lymphoma in XLP-2 patients.

**Key points:**

1. XIAP loss-of-function markedly impairs EBV+ B-cells outgrowth over the first week post-infection, particularly in the presence of IFN-γ.
2. XIAP mutation impedes EBV-driven B-cell transformation by potentiating p53-driven caspase activation and apoptosis.

## Introduction

EBV persistently infects more than 90% of the global population and contributes to over 200,000 cancers per year ^1–3^. While acute EBV infection is controlled by immunocompetent hosts and typically results in subclinical disease, EBV is nonetheless a major driver of pathology in hosts with primary or acquired immunodeficiency, in particular lymphoproliferative disease ^4–8^. EBV is associated with 200,000 cancers per year, including lymphomas, gastric and nasopharyngeal carcinoma ^9^. Defects in anti-EBV immunity increase rates of multiple EBV-associated lymphomas including Burkitt, Hodgkin and immunoblastic lymphomas ^5,7,10^.

Immune control over EBV is primarily mediated by T-cell responses, particularly CD8^+^ cytotoxic T lymphocytes, which target virus-infected cells. Natural killer (NK) cells and antibodies also play crucial roles in regulating EBV activity^11^. The importance of these immune mechanisms is underscored by the severe outcomes observed in individuals with compromised immunity, such as those with X-linked lymphoproliferative syndromes (XLP)^11^. In these cases, defects in immune signaling pathways can lead to uncontrolled viral proliferation and associated complications, highlighting the critical balance between EBV and host immune responses.

Two rare congenital XLP syndromes have been described, which share in common extreme susceptibility to EBV, frequently resulting in fulminant infectious mononucleosis, dysgammaglobulinemia and hemophagocytic lymphohistiocytosis (HLH). HLH is a T-cell and macrophage hyperactivation state ^12–14^. XLP-1 is caused by loss-of-function mutations in *SH2D1A*, which encodes the protein 128 amino acid SH2-domain containing signaling lymphocyte activation molecule (SLAM)–associated protein (SAP) (MIM no. 308240). SAP controls signaling downstream of SLAM family receptors, including CD150, CD229, 2B4, CD84 and NTB-A ^15,16^. XLP-2 instead arises from congenital mutations of the X-linked inhibitor of apoptosis (XIAP, also termed BIRC4; MIM no. 300635), a 497-amino acid member of the inhibitor of apoptosis protein (IAP) family that serves as a central regulator of apoptotic cell death by inhibiting caspases 3 and 7 ^15–19^. XIAP has additional roles in many other pathways ^20^. Each XLP syndrome is characterized by markedly elevated EBV viral loads ^21,22^, which trigger severe infectious mononucleosis, HLH and a range of hematological dyscrasias. While rates of HLH and splenomegaly are higher with XLP-2, there are no reported cases of EBV-associated lymphoproliferative disease in this syndrome. This stands in contradistinction to essentially all other primary immunodeficiency syndromes manifest by susceptibility to EBV ^21,23^. Much remains to be learned about why *SH2D1A* and *XIAP* mutations cause XLP syndromes, but each is associated with defects in T and NK cell responses, including the absence of natural killer T-cells ^7^.

A notable difference between SAP and XIAP is their tropism. While SAP is expressed primarily in NK, NKT and T cells, XIAP is ubiquitously expressed in all cell types ^21^. This disparity suggests that the lack of B-cell malignancies in XLP-2 may be attributed to intrinsic factors within EBV-infected B-cells themselves, rather than solely to differences in immune cell function. Gene expression analysis highlighted elevated levels of the tumor suppressor cell adhesion molecule 1 (CADM1) in XIAP-deficient B-cells infected by EBV ^24^, which has been implicated in NF-kB activation in B-cells infected by the gamma-herpesvirus Kaposi’s Sarcoma Associated Herpesvirus ^25^.

Here, we sought to test the role of XIAP in EBV-driven lymphomagenesis. We found that the knockout or mutation of *XIAP* in primary B-cells impacted the EBV-driven proliferation of newly infected B-cells, leading to reduced efficiency in B-cell transformation. We provide evidence that XIAP supports newly EBV-infected primary B-cell survival, which counteracted upregulation of p53-related apoptosis signaling triggered by EBV infection. B-cell intrinsic XIAP deficiency elevated the apoptosis frequency at early timepoints of EBV-mediated B-cell immortalization but not in cells stimulated by CD40-ligand (CD40L) and interleukin-4 (IL-4), exacerbated by the presence of inflammatory cytokines.

## Materials and methods

### Cell cultures and Chemical compounds

Cells were cultured following the vendor’s instructions. Details are in the supplemental materials and methods. Chemical compounds used in the study are listed in the supplemental materials and methods.

### Primary Human B cells

Leukocyte fractions that were discarded and de-identified, originating from platelet donations, were obtained from the Brigham and Women’s Hospital Blood Bank. These fractions were utilized for the isolation of primary human B cells following our Institutional Review Board-approved protocol. Venous blood of XLP2 patients and corresponding controls were obtained from Boston Children’s Hospital. PBMCs were isolated using Lymphopre Density Gradient Medium (Stem Cell Technologies), and primary B cells were subsequently isolated by negative selection using RosetteSep Human B Cell Enrichment and EasySep Human B cell enrichment kits (Stem Cell Technologies), according to the manufacturers’ protocols. Cells were cultured in RPMI-1640 medium with 10% FBS.

### EBV production

For production of Akata EBV, Akata EBV+ cells were resuspended in FBS-free RPMI-1640 at a concentration of 2-3×10^6^ cells per ml and treated by 0.3% Polyclonal Rabbit Anti-Human IgG (Agilent) for 6 hours. Cells were cultured in RPMI-1640 with 4% FBS for 3 more days, and the virus-containing supernatants were collected by ultracentrifugation and filtration through a 0.45 μm filter. The viral titer was determined by EBV transformation assay as described below.

### EBV transformation assay

Purified human primary B cells were seeded into a 96 well plate at 50000 cells per well. The stock of Akata EBV was diluted ten-fold in order, and 100 μL of virus dilution was added to each well. The cells were maintained in RPMI-1640 with 10% FBS at 37°C. After 4 weeks of incubation, the proportion of wells with B cell outgrowth was scored. A transforming unit per well was defined as the virus quantity necessary for achieving B cell outgrowth in 50% of wells. The multiple of infection (MOI) was determined by dividing the Transforming Unit by the cell number.

### CRISPR/Cas9 editing

For cell lines with stable Cas9 expression, sgRNA sequences from Broad Institute Brunello library were used. sgRNA oligos were cloned into the pLentiGuide-Puro vector (Addgene plasmid #52963, a gift from Feng Zhang), and used for lentivirus production in HEK293T cells. After 2 rounds of transduction performed at 48 and 72 hours post plasmids transfection, cells were selected by 3 μg/ml puromycin for more than 4 days.

For CRISPR/Cas9 editing in primary B cells, Cas9 RNA complexes were transduced into the cells with electroporation. In brief, crRNA targeting XIAP was selected using Alt-R Predesigned Cas9 crRNA Selection Tool from Integrated DNA Technologies. TracrRNA and Cas9 Nuclease V3 were also obtained from Integrated DNA Technologies. The crRNA and tracrRNA were annealed to form the duplex and incubated with Cas9 for 20 minutes. Then the cells were mixed with the RNP complexes, and electroporated using the Neon NxT Electroporation System at 1700V, 20ms and 2 pulses. sgRNA sequences were listed in Table S1.

## Results

### XIAP supports the outgrowth of newly EBV-infected primary human B-cells

EBV transforms primary human B-cells into immortalized, continuously proliferating lymphoblastoid cell lines (LCLs), which serves as a major model for EBV lymphomagenesis. To characterize how XIAP deficiency affects EBV-mediated B-cell immortalization, we utilized CRISPR-Cas9 technology to knockout (KO) *XIAP* in primary B-cells isolated from healthy donors (Figure 1A). FACS analysis indicated that Cas9/single guide RNA (sgRNA) ribonucleoprotein complexes (RNP) were successfully delivered to >50% of B-cells (Figure S1A), and immunoblot analysis confirmed depletion of XIAP expression across the bulk population (Figure 1B).

**Figure 1.**
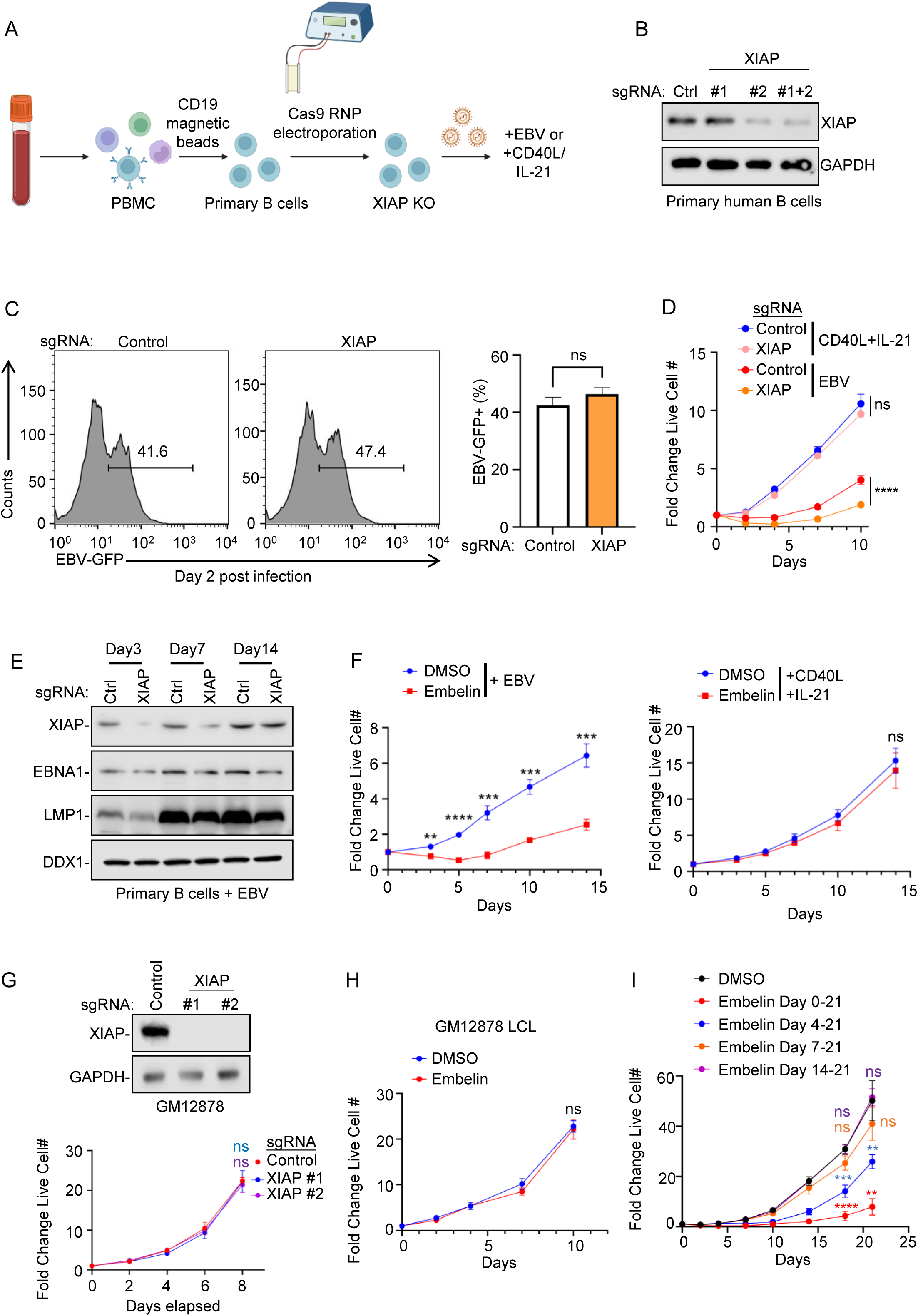
XIAP inactivation impairs the outgrowth of newly EBV infected primary B-cells. (A) Workflow for electroporation and EBV infection of primary human B-cells. B-cells purified from peripheral blood mononuclear cells (PBMC) were transduced with Cas9 ribonucleoprotein (RNP) complexes containing *XIAP* targeting or non-targeting control single guide RNA (sgRNA). 1 hour post electroporation, cells were infected with EBV or stimulated by CD40 ligand (CD40L) (50 ng/ml) and IL-21 (50 ng/ml). (B) Immunoblot analysis of whole cell lysates (WCL) from primary B-cells on day 3 post electroporation with Cas9 control (ctrl) or XIAP sgRNA containing RNPs. (C) XIAP editing does not alter EBV infection efficiency. FACS analysis of control versus XIAP edited B-cells at Day 2 post-infection by recombinant EBV that expresses a green fluorescence protein (GFP) marker. Mean + standard deviation (SD) GFP+ cell percentages from n=3 replicates are shown on the right. (D) Growth curve analysis of primary human B-cells transfected with the indicated sgRNA-containing Cas9 RNPs and treated with CD40L and IL-21 or infected with EBV. CD40L/IL-21 were replenished every 3 days. Mean ± SD fold change live cell numbers from n = 3 biological replicates, relative to Day 0 values, are shown. (E) Immunoblot analysis of WCL from primary B-cells with control or XIAP sgRNA-containing RNP on the indicated days post-EBV infection. (F) Growth curve analysis of primary B-cells treated with DMSO or the XIAP inhibitor embelin (5 μM) together with EBV infection (left) or CD40L/IL-21 treatment (right). Embelin, CD40L and IL-21 were replenished every 3 days. Mean ± SD fold change live cell numbers from n=3 replicates, relative to Day 0 values, are shown. (G) Immunoblot and growth curve analysis of Cas9+ GM12878 LCLs expressing the indicated control or independent *XIAP* targeting sgRNAs. Mean ± SD fold change live cell numbers from n=3 replicates, relative to Day 0 values, are shown. (H) Growth curves analysis of GM12878 LCLs treated with DMSO or embelin (5 μM), which were replenished every 3 days. Mean ± SD fold change live cell numbers from n=3 replicates, relative to Day 0 values, are shown. (I) Growth curve analysis of primary human B-cells infected by EBV at Day 0 and then treated with embelin (5 μM) over the indicated times. Embelin was replenished every 3 days. Mean ± SD fold change live cell numbers from n=3 replicates, relative to Day 0 values, are shown. Statistical significance was assessed by comparing each indicated groups with DMSO control groups. Statistical significance was assessed by two-tailed unpaired Student’s t test (C, D, F, H, I) or one-way ANOVA followed by Tukey’s multiple comparisons test (G). Blots are representative of n=3 replicates. **p<0.01, ***p<0.001, ****p<0.0001, ns, not significant.

Control versus XIAP depleted B-cells were infected with EBV, or for cross-comparison, stimulated by CD40L and IL-21, a combination which efficiently drives B-cell proliferation ^26^. *XIAP* editing did not significantly alter EBV infection efficiency, as judged by an EBV genomic green fluorescence protein (GFP) reporter that can be used to mark infected B-cells (Figure 1C). However, growth curve analysis highlighted that *XIAP* editing markedly reduced the efficiency of EBV-driven primary B-cell outgrowth. Intriguingly, XIAP KO did not significantly alter proliferation of CD40L/IL-21 treated cells, suggesting an EBV-specific, B-cell intrinsic phenotype (Figure 1D). Our CRISPR editing only depleted XIAP in a subset of B-cells, likely due to the inability to deliver Cas9 RNPs across the population. Consistent with a growth advantage for XIAP-expressing cells, immunoblot analysis highlighted that there was a selection against XIAP deficiency evident by day 14 post-infection, with similar XIAP levels present in control vs XIAP edited populations, in contrast to earlier timepoints (Figure 1E).

To further characterize a potential XIAP role in support of early EBV-mediated B-cell outgrowth, we treated newly infected versus CD40L/IL-21 stimulated B-cells with the small-molecule XIAP antagonist embelin ^27^. Consistent with the *XIAP* KO phenotype, embelin significantly impeded EBV-driven but not CD40L/IL21-induced B-cell outgrowth (Figure 1F). Taken together, these results indicate that XIAP plays a B-cell intrinsic role in EBV, but not CD40L/IL-21 driven proliferation. By contrast, consistent with prior reports ^24^, XIAP KO or inhibition by embelin in established LCLs failed to significantly alter proliferation of either GM12878 or GM15892 LCLs, despite excellent efficiency of CRISPR editing (Figure 1G-H, S1B-C), suggesting that XIAP may specifically play a critical role at an early stage of EBV-mediated B-cell transformation. In support of this hypothesis, early administration of embelin impaired EBV-driven outgrowth, but its impact was non-significant when started at 7 days post-EBV infection or at later timepoints (Figure 1I). Collectively, these findings suggest that XIAP plays a critical role in support of EBV-infected B-cell transformation within the first week of infection.

### Newly EBV-infected but not CD40L/IL-21 driven proliferation is impaired in XLP-2 B-cells

We next characterized effects of XIAP deficiency on early EBV-mediated B-cell outgrowth in primary B-cells from XLP-2 patients versus from healthy controls. XLP-2 patients 1 and 2, who are brothers, possess an XIAP missense variant (NP_001158.2: p.Ser421Asn) that compromises XIAP function (Figure 2A) ^28^. Venous blood was collected from three healthy donors on the same day (Control 1-3). B-cells were purified by negative selection and infected by EBV or stimulated by CD40L/IL-21. Similar to our XIAP CRISPR analyses, proliferation of XLP-2 cells was diminished over the first week of infection relative to healthy controls (Figure 2B). By contrast, XLP-2 and control B-cells proliferated similarly in response to CD40L and IL21-treatment (Figure 2C). We observed similar effects with B-cells from two additional XLP-2 patients (Patients 3 and 4) who possess an XIAP nonsense variant (NP_001158.2: p.Arg49*) (Figure S2A-C). Consistent with a key early but not late XIAP role in support of EBV-mediated transformation, LCLs established from both XLP-2 patients and from controls proliferated at similar rates (Figure 2D).

**Figure 2.**
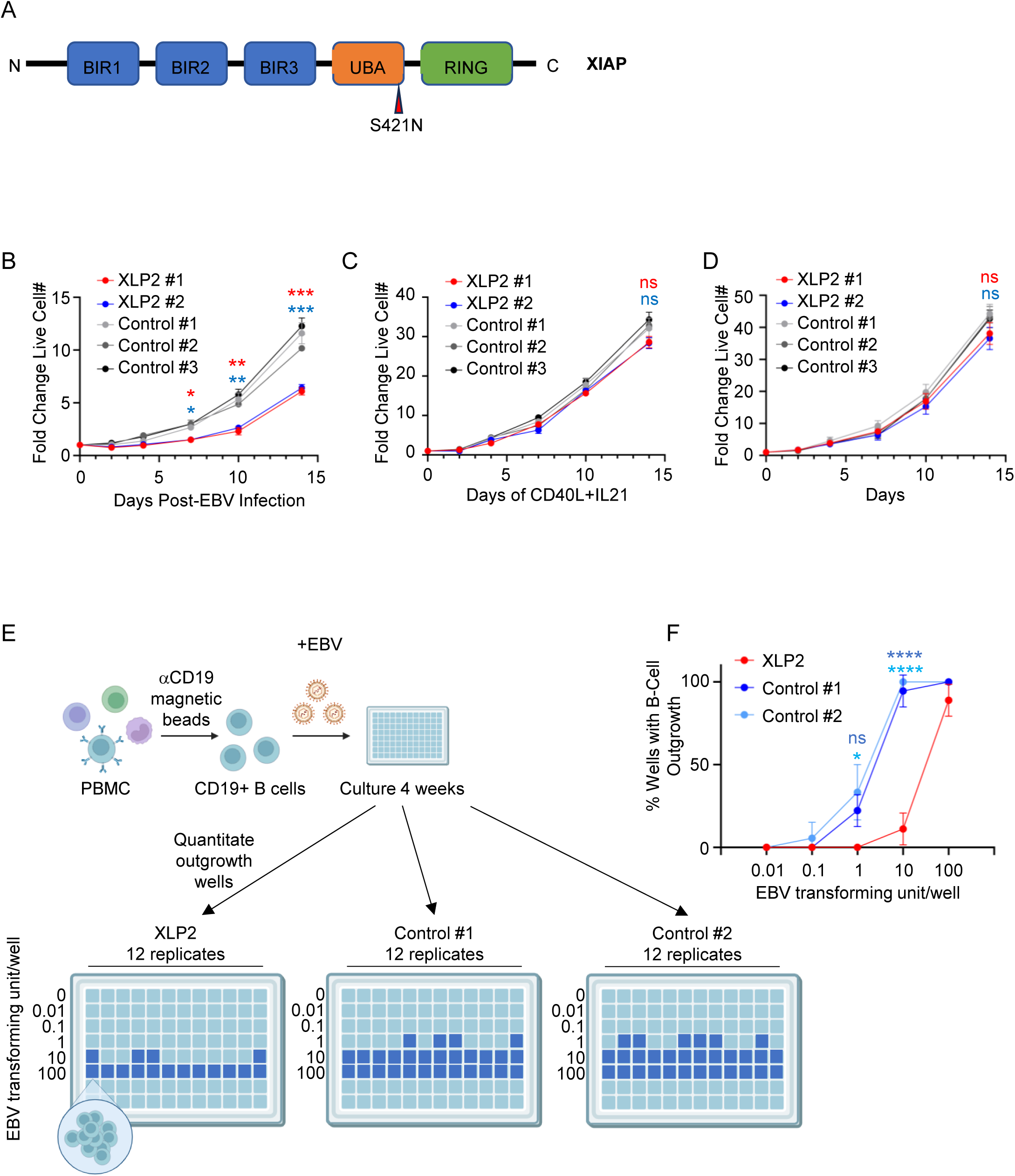
XLP2 patient B-cells demonstrate impaired EBV but not CD40L/IL-21 driven outgrowth at early timepoints. (A) Schematic diagram highlighting the *XIAP* mutation shared by XLP2 Patients #1 and #2. (B) Growth curve analysis of primary B-cells from XLP2 patients or from controls that were infected with EBV. Mean ± SD fold change live cell numbers from n=3 replicates, relative to Day 0 values, are shown. The annotations represent the results of statistical comparisons between XLP2 samples and Control #1. (C) Growth curve analysis of primary B-cells from XLP2 patients or controls treated with CD40L and IL-21, which was replenished every 3 days. Mean ± SD fold change live cell numbers from n=3 replicates, relative to Day 0 values, are shown. The annotations represent the results of statistical comparisons between XLP2 samples and Control #1. (D) Growth curves of lymphoblastoid cells established from B cells from either two XLP2 patients or three controls. Mean ± SD fold change live cell numbers from n=3 replicates, relative to Day 0 values, are shown. The annotations represent the results of statistical comparisons between XLP2 samples and Control #1. (E) EBV B-cell transformation assay workflow. CD19+ B-cells purified from PBMCs were plated and infected with serial dilutions of the Akata EBV strain, using a range of 0 – 100 EBV transforming units/well. Wells with B-cell outgrowth were scored 4 weeks later. Outgrowth wells on one of the three replicate plates were displayed. (F) EBV transformation assays of primary human B-cells from XLP2 patient or healthy controls, as in (E). Shown are the mean ± SD percentages of wells with B-cell outgrowth from n=3 replicates. Statistical significance was assessed by one-way ANOVA followed by Tukey’s multiple comparisons test (B-D, F). *p<0.05, **p<0.01, ***p<0.001, ****, p<0.0001, ns, non-significant.

To further characterize *XLP-2* mutation effects, we conducted transformation assays, in which serial dilutions of EBV are added to primary B-cells, and the percentage of wells with cellular outgrowth are scored at 4 weeks post-infection (Figure 2E). Consistent with our growth curve phenotypes, *XIAP* mutation significantly reduced EBV B-cell transformation efficiency (Figure 2F). Together, these findings demonstrate the limited extent of EBV-induced B-cell transformation in XLP2 patients.

### XIAP plays key anti-apoptosis roles in newly-EBV infected B-cells

To explore the mechanism by which XIAP supports EBV but not CD40/IL-21-driven B-cell outgrowth, we tested the effects of XIAP depletion on growth versus survival at early times post-EBV infection. Interestingly, CRISPR XIAP editing significantly impaired proliferation of EBV infected but not CD40L/IL-21 stimulated cells (Figure 3A-3B). Furthermore, FACS analysis of 7-AAD vital dye uptake revealed an increased percentage of cell death in XIAP edited and EBV-infected, but not CD40L/IL-21 stimulated B-cells (Figure 3C).

**Figure 3.**
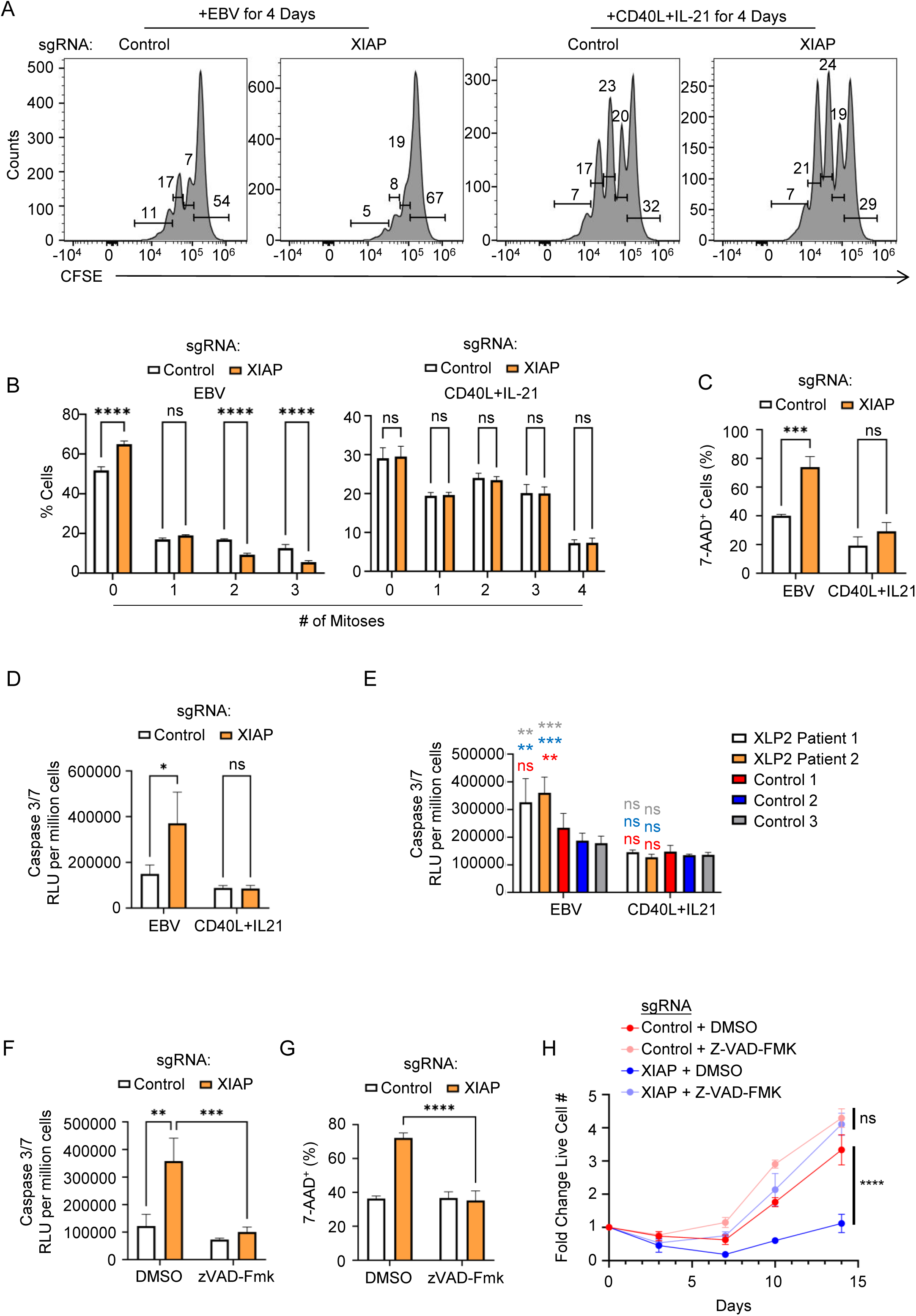
EBV but not CD40L/IL-21 triggers apoptosis within the first week of XLP2 B-cell infection. (A) FACS analysis of control versus XIAP edited primary human B-cells at Day 4 post-EBV infection or CD40L/IL-21 stimulation. Shown are representative FACS plots from n=3 replicates of cells stained with CFSE prior to EBV infection or CD40L/IL-21 treatment. Live cells were gated by absence of 7-AAD vital dye uptake. (B) Mean + SD Percentages of cells with the indicated number of mitoses from n=3 replicates as in (A) of EBV infection versus CD40L/IL-21 stimulation. (C) Mean + SD % 7AAD+ cells from n=3 replicates of control or XIAP edited primary B-cells on Day 4 post-EBV infection or CD40L/IL21 treatment. (D) Mean + SD caspase 3/7 activity from n=3 replicates of control or XIAP edited primary B-cells on day 4 post-EBV infection or CD40L/IL-21 treatment. (E) Mean + SD caspase 3/7 activity from n=3 replicates of control or XLP2 primary B-cells on day 4 post-EBV infection or CD40L/IL-21 treatment. (F) Mean + SD caspase 3/7 activity from n=3 replicates of control or XIAP edited primary B-cells incubated with DMSO vehicle or the pan-caspase inhibitor zVAD-fmk (20 μM) on day 4 post-EBV infection or CD40L/IL-21 treatment. (G) Mean + SD % 7AAD+ cells from n=3 replicates of control or XIAP edited primary B-cells on Day 4 post-EBV infection or CD40L/IL21 treatment. (H) Growth curve analysis of control versus *XIAP* edited primary B-cells infected with EBV on Day 0 and cultured with DMSO vehicle or zVAD-Fmk (20 μM). Mean ± SD fold change live cell numbers, relative to uninfected values, are shown. DMSO or zVAD-Fmk were replenished every 3 days. Statistical significance was assessed by two-tailed unpaired Student’s t test (B-D, H) or one (E) or two-way (F and G) ANOVA followed by Tukey’s multiple comparisons test. In A-G, CD40L and IL21 were replenished on Day 3. **p<0.01, ***p<0.001, ****p<0.0001. ns, not significant.

We hypothesized that XIAP deficiency might sensitize EBV-infected cells to apoptotic cell death, given XIAP’s ability to block executioner caspase activity, including caspases 3 and 7 ^29^. In support, caspase 3 and 7 activity was significantly elevated in XIAP edited cells on day 4 post-EBV infection but not at the same timepoint of CD40L/IL-21 stimulation (Figure 3D). Caspase 3/7 activity was similarly elevated at Day 4 post-EBV-infection but not CD40L/IL-21 stimulation of XLP-2 patient primary B-cells (Figure 3E). We therefore tested whether caspase activity was necessary for EBV-triggered cell death of XIAP-deficient B-cells. The pan-caspase inhibitor zVAD-Fmk significantly inhibited caspase 3/7 activity and blocked EBV-driven cell death of XIAP edited cells (Figure 3F-G). zVAD-Fmk also significantly increased the outgrowth of EBV-infected *XIAP* CRISPR edited cells (Figure 3H). These results suggest that EBV infection drives an apoptotic stimulus within the first week of primary B-cell infection, a period in which EBV drives Burkitt-like hyperproliferation ^30–32^, and that XIAP anti-caspase activity is critical to overcome EBV-triggered caspase activity in support viral mediated B-cell transformation.

### EBV activates p53 induced apoptosis signaling

We next aimed to decipher the mechanism behind XIAP’s differential impact on EBV-infected but not CD40L/IL-21 proliferation. To gain insights, we performed systematic transcriptomic and whole cell proteomic analyses of XLP-2 versus healthy control B-cells at Day 7 post-EBV infection or CD40L/IL-21 stimulation. This analysis highlighted that EBV infection upregulated expression of P53 (encoded by *TP53*) relative to levels in CD40L/IL-21 stimulated cells. EBV also altered expression levels of multiple apoptosis pathway components, increasing levels of the pro-apoptotic BAX, NOXA (encoded by PMAIP1) and PUMA (encoded by BBC3) in both control and XLP2 patient B-cells (Figure 4A and 4B and Table S2 and S3). Intriguingly, many of these genes are associated with the p53 signaling pathway. Notably, these data are consistent with prior transcriptomic and proteomic analyses ^30,33^ of peripheral blood B-cell EBV infection, which identified that levels of TP53 and BAX peak at day 4 post-EBV infection and then gradually decline (Figure 4C).

**Figure 4.**
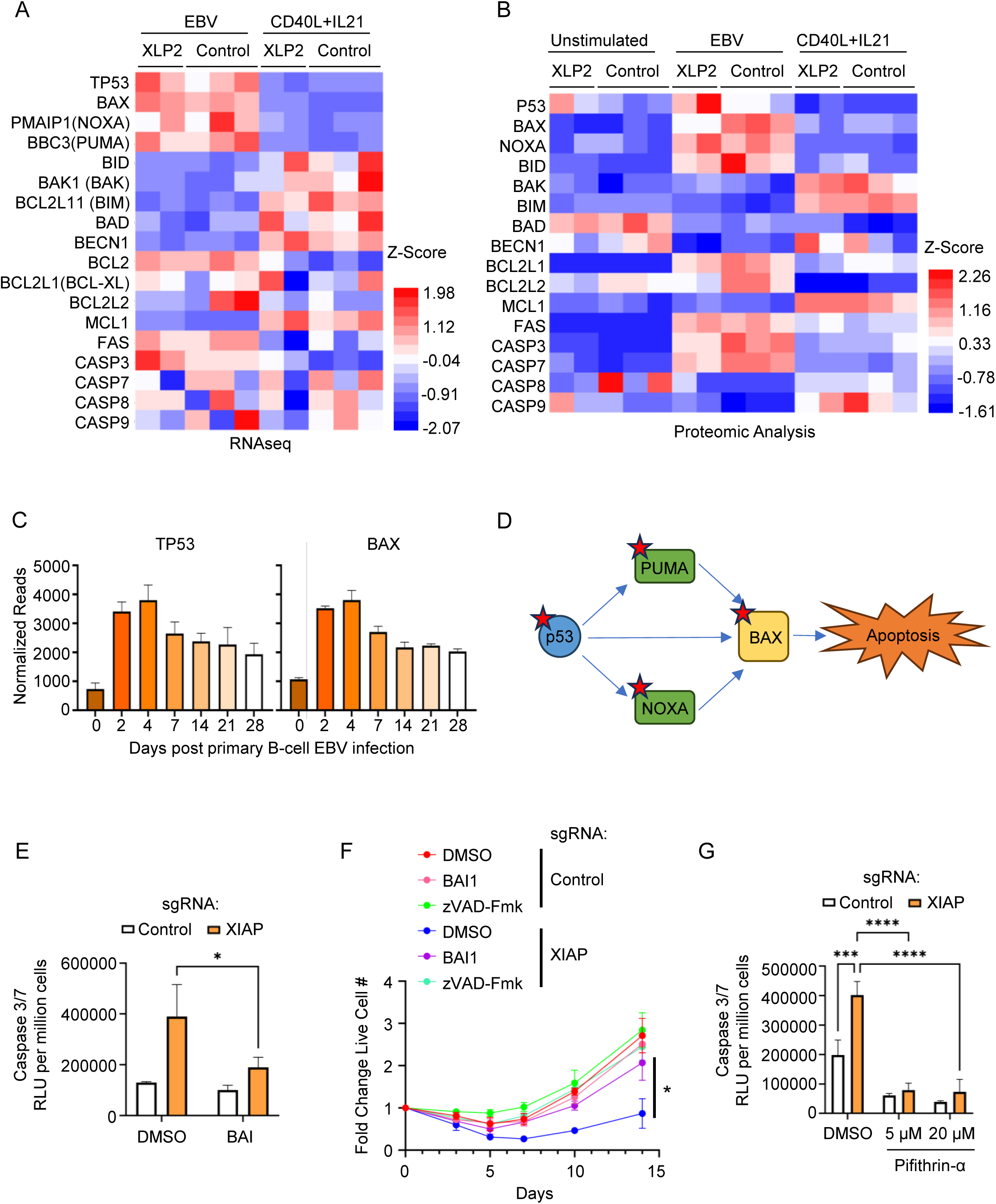
EBV but not CD40L/IL-21 activates p53- and BAX-dependent apoptosis in newly infected XIAP deficient B-cells. (A) Heatmap of RNAseq analysis of primary B-cells from XLP2 patient or controls on Day 7-post EBV infection or CD40L/IL-21 treatment. Z-scores of normalized reads of the indicated mRNAs from n=3 replicates are shown. Columns show data from two XLP2 patients or three controls. (B) Multiplexed tandem mass tag proteomic analysis of primary B-cells from XLP2 patients or healthy controls on Day 7 post-EBV infection or CD40L/IL-21 treatment. Unstimulated cells were harvested on Day 0. Z-scores of relative protein abundances of the indicated mRNAs from n=3 replicates are shown. Columns show data from two XLP2 patients or three controls. (C) Mean + standard deviation (SD) TP53 and BAX mRNA expression from n=3 replicates of RNAseq analysis of primary human B-cells on the indicated days post EBV infection ^33^. (D) Schematic diagram illustrating p53 that p53 target genes PUMA and NOXA can each upregulate BAX, which in turn induces the intrinsic apoptosis pathway. Red stars denote upregulation at Day 7 post-EBV infection relative to CD40L/IL-21 levels. (E) Mean + SD caspase 3/7 activity from n=3 replicates of primary B-cells expressing control or XIAP targeting sgRNAs and treated with BAI1 (5 μM) on day 4 post EBV infection. BAI1 was added from Day 0 onwards, replenished every 3 days. (F) Growth curve analysis of control versus *XIAP* edited primary B-cells and cultured with DMSO, zVAD-Fmk (20 μM) or BAI1 (5 μM) from Day 0 onwards. Mean ± SD fold change live cell numbers, relative to uninfected values, are shown. DMSO, BAI and zVAD-Fmk were replenished every 3 days. (G) Mean + SD caspase 3/7 activity from n=3 replicates of control versus *XIAP* edited primary B-cells cultured with DMSO or pifithrin-α at 5 or 20 μM on Day 4 post-infection. Statistical significance was assessed by two-tailed unpaired Student’s t test (F) or two-way ANOVA followed by Tukey’s multiple comparisons test (E and G). *p<0.05, ***p<0.001, ****p<0.0001.

P53, as well as multiple p53-upregulated pro-apoptotic proteins can induce expression of the pro-apoptosis BCL-2 family member BAX ^34,35^ (Figure 4D). We therefore tested whether the small molecule allosteric BAX inhibitor BAI1 ^36^ could suppress apoptosis induction in newly EBV-infected XIAP deficient cells. Interestingly, BAX blockade by BAI significantly diminished EBV-driven caspase 3/7 activity in XIAP deficient cells (Figure 4E). Similarly, BAI significantly restored EBV-mediated outgrowth of CRISPR XIAP-edited B-cells (Figure 4F). In further support of a key p53 role, p53 inhibition by pifithrin-α ^37^ also significantly inhibited EBV-driven caspase 3/7activity in XIAP depleted primary B-cells (Figure 4G). These results suggest that EBV upregulates p53-driven BAX activation, whose activation of downstream caspase signaling necessitates XIAP to inhibit apoptosis.

### XIAP in EBV transformed cells renders sensitivity to inflammatory cytokines

While the above experiments indicate a role for XIAP in protecting newly EBV infected cells from apoptosis, a subset of EBV+ B-cells survive and can be immortalized, raising the question of why this does not apparently lead to lymphoma in XLP-2 patients. One hypothesis is, other factors in vivo may further influence the transformation of EBV-infected XIAP deficient cells. To test this hypothesis, we first treated peripheral blood mononuclear cells (PBMC) from healthy donors with the small-molecule XIAP antagonist embelin, which interacts with the same XIAP BIR3 domain residues as caspase-9 ^27^, to mimic the in vivo EBV infection of XLP2 patients.

Embelin treatment significantly reduced the percentage of CD19+ B-cells at 7-day post-infection, a timepoint at which most surviving B-cells are EBV-infected. However, embelin did not significantly reduce CD56+ NK, CD4+ or CD8+ T-cell frequency in PBMC cultures (Figure 5A-B), suggesting an XIAP role in infected B-cell survival. To exclude the possibility that embelin exhibits B cell-specific toxicity, we treated primary B cells with embelin prior to either EBV infection or CD40L/IL21 stimulation. Consistent with our XIAP knockout results, embelin-treated cells showed increased cell death only in EBV-infected B cells, while CD40L/IL21-stimulated cells remained unaffected (Figure 5C). These findings suggest that embelin’s effects closely mimic those observed in XIAP-deficient cells, specifically targeting EBV-infected B cells without general B cell toxicity.

**Figure 5.**
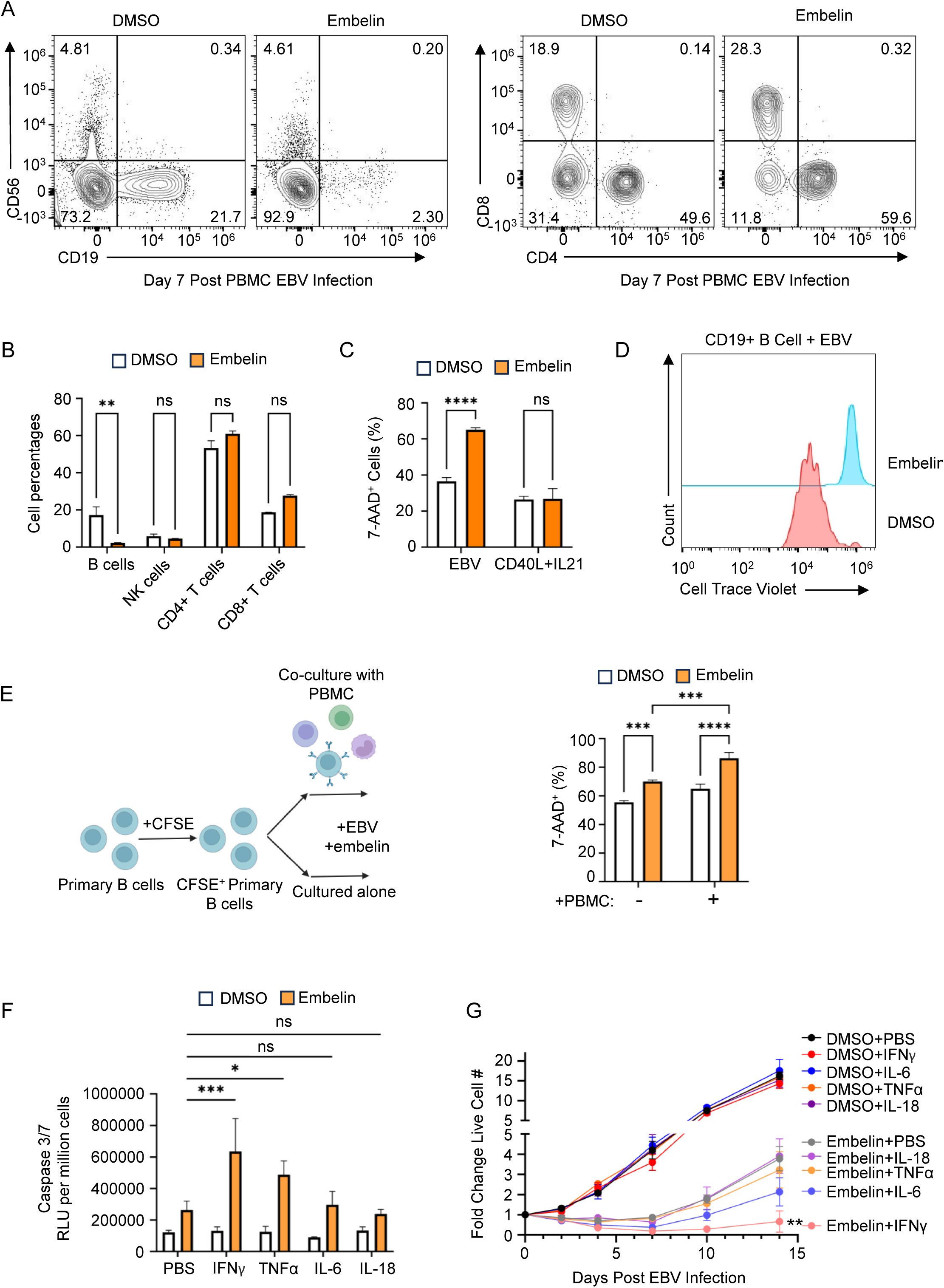
Embelin XIAP inhibition perturbs EBV-mediated primary B-cell outgrowth and sensitizes newly-infected cells to IFNγ-triggered apoptosis. (A) FACS analysis of CD4+ or CD8+ T cell, CD56^+^ NK cell and CD19+ B-cell subsets from PBMCs of a control donor, infected with EBV and treated with DMSO or embelin (5 μM) on Day 7 post-EBV infection. DMSO and embelin were added starting from Day 0 and replenished every 3 days. (B) Mean + SD percentages of CD4+ or CD8+ T cell, CD56^+^ NK cell and CD19+ B-cells from (A) are shown. (C) Mean + SD % 7AAD+ cells from n=3 replicates of DMSO or embelin treated primary B-cells on Day 4 post-EBV infection or CD40L/IL21 treatment. DMSO and embelin were added starting from Day 0 and replenished every 3 days. (D) FACS analysis of CD19+ B-cells CellTrace Violet (CTV) dye dilution, whose abundance is reduced by half with each mitosis, from PBMCs treated with DMSO (control) or embelin (5 μM) on Day 7 post-EBV infection. The whole PBMC cell cultures were stained with CTV and infected with EBV and treated with DMSO or embelin (5 μM) as in (A). Cells were stained with anti-CD19 antibodies on day 7 and CTV on CD19+ B cell was analyzed. DMSO and embelin were added starting from Day 0 and replenished every 3 days. (E) Mean + SD percentages of 7AAD+ cells of primary human B-cells from controls, cultured alone or co-cultured with autologous PBMC in the presence of DMSO versus embelin (5 μM) for 4 days. B-cells were stained with CFSE prior to PBMC co-culture, which served as a cell trace marker to allow their FACS gating within mixed PBMC cultures. (F) Mean + SD caspase 3/7 activity from n=3 replicates of EBV-infected DMSO or embelin treated cells that were co-cultured with PBS vehicle, IFNg (50 ng/mL), TNFa (50 ng/mL), IL-6 (50 ng/mL) or IL-18 (50 ng/mL) on Day 4 post-infection. (G) Growth curve analysis of control primary human B-cells infected by EBV and treated with DMSO or embelin (5 μM) together with IFNγ, TNFα, IL-6 or IL-18 as indicated. Shown are mean ± SD fold change live cell numbers from n=3 replicates. Statistical significance was assessed by comparing each cytokine treated group with PBS control group. Statistical significance was assessed by two-tailed unpaired Student’s t test (B, C, G) or two-way ANOVA followed by Tukey’s multiple comparisons test (E and F). In all experiments, DMSO, Embelin and cytokines were refreshed every 3 days. *p<0.05, **p<0.01, ***p<0.001, ****p<0.0001. ns, not significant.

Embelin reduction of EBV-infected B-cell numbers could be by effects on growth and/or survival. To investigate this further, we stained the PBMC with CellTrace Violet and analyzed the CD19+ B cells with FACS dye-dilution analysis. Interestingly, we observed that markedly less proliferation of EBV-infected CD19+ B-cells in embelin treated PBMC cultures, though analysis was complicated by low numbers of live B-cells at this timepoint in embelin treated PBMCs (Figure 5D). We therefore next analyzed embelin effects on CD19+ cell death, as judged by 7-AAD uptake. Interestingly, embelin XIAP blockade increased EBV-infected B-cell cell death when cultured with autologous PBMCs than when cultured alone (Figure 5E). These results thus suggest that the presence of other immune cells further enhances the apoptosis of EBV-infected cells.

We next sought to determine the factor that promotes the apoptosis of EBV infected cells. Notably, multiple pro-inflammatory cytokine levels are elevated in XLP-2 patient serum in particular IL-18, IL-6, interferon gamma (IFNγ) and tumor necrosis factor alpha (TNFα) ^38^. EBV infection also upregulates infected B-cell expression of multiple pro-inflammatory cytokines, and IL-18, TNFα and IFNγ mRNAs are upregulated within the first week of EBV infection ^33^. Since pro-inflammatory cytokines can sensitize cells to apoptosis, we hypothesized that XIAP deficiency and pro-inflammatory cytokines might exert synthetic effects that together potentiate apoptosis of newly EBV-infected cells. To test this hypothesis, we treated EBV-infected purified B-cells from three healthy donors with vehicle or embelin, together with vehicle, IFNγ, TNFα, IL-5 or IL-18. Intriguingly, IFNγ and to a lesser extent TNFα treatment significantly increased caspase 3/7 activity in embelin but not DMSO-treated cells (Figure 5F). Interestingly, IFNg treatment also strongly suppressed outgrowth of embelin treated but not vehicle treated EBV-infected B-cells over the first two weeks (Figure 5G). These findings indicate that XIAP inhibition in newly EBV-infected cells renders the cells sensitive to inflammatory cytokines.

Collectively, our data is consistent with a model in which EBV triggers p53-mediated apoptotic signaling, which results in activation of caspase 3 and 7. Existence of XIAP in the host cells can inhibit this signaling, thus allowing for cell survival and B cell transformation. In patients with XLP2, mutation of XIAP makes the cells sensitive to p53-mediated apoptotic signaling and inflammatory cytokines, resulting in cell death of EBV infected cells, thus abolishing lymphomagenesis (Figure 6).

**Figure 6.**
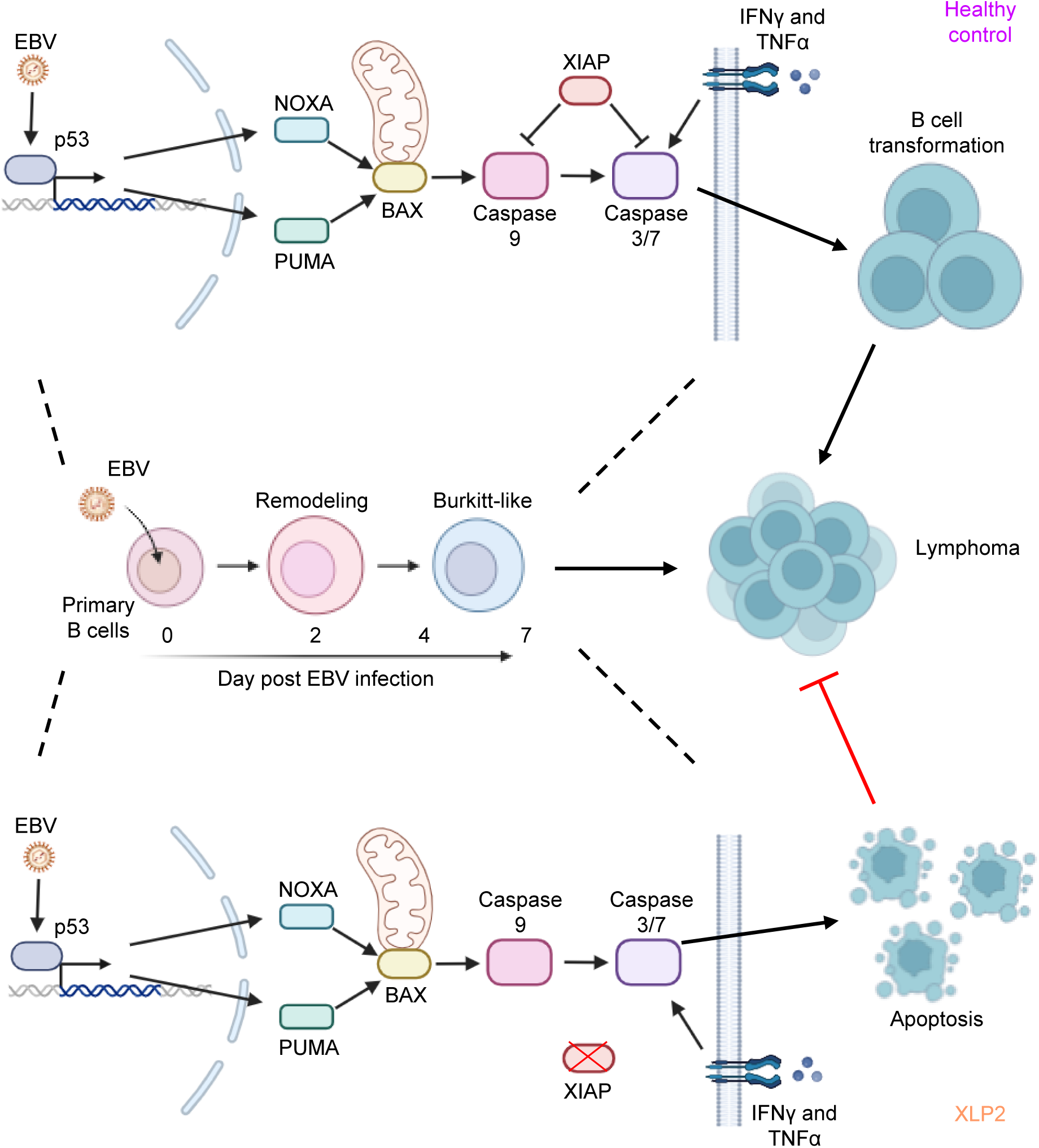
Schematic model of key anti-apoptotic XIAP role in newly EBV-infected B-cells. EBV drives rapid proliferation of newly infected B-cells, which triggers DNA damage, upregulation of p53 and of downstream NOXA, PUMA and BAX. XIAP blocks caspase activity and apoptosis in most settings, including with XLP1, enabling newly EBV-infected B-cells to undergo transformation and in XLP1 to cause high rates of lymphomas. Lymphomas are not observed in XLP2 patients, where the absence of XIAP enables caspase 3/7 activation and apoptosis induction over the first week of EBV infection, which is exacerbated by the inflammatory cytokine milieu, in particular IFNγ, restraining lymphomagenesis.

## Discussion

The absence of EBV-associated lymphoma is a striking feature of XLP2, which separates it from XLP1 and from nearly all other immunodeficiencies with defective cell mediated control of EBV-infected B-cells. Here, we used multidisciplinary approaches to characterize how XIAP perturbation by XLP2 mutation, CRISPR editing or small molecule inhibition alters EBV/B-cell interactions and early events in EBV-driven B-cell transformation. Multi-omic profiling highlighted that in the absence of XIAP, EBV upregulates caspase 3 and 7 activity in a p53 and BAX dependent manner over the first week of primary B-cell infection, limiting transforming B-cell outgrowth. Co-incubation with IFNγ or TNFα, which are found in XLP2 serum at elevated levels in particular in the setting of HLH, strongly further suppressed EBV immortalization of XIAP depleted B-cells.

XLP2 lymphocytes display increased susceptibility to extrinsic apoptosis triggered by CD95/FAS, TNFα or TNF related apoptosis-inducing ligand (TRAIL), and XLP2 T-cells also exhibit increased rates of activation-induced programmed cell death ^18,39^. However, EBV can convert XLP2 B-cells with loss-of-function XIAP mutations into continuously growing, immortalized lymphoblastoid cell lines (LCLs) in culture ^24^, which are a key model for EBV-driven immunoblastic lymphomas of immunocompromised hosts. Our findings indicate that XIAP is not critical for growth or survival of infected cells that convert to lymphoblastoid physiology over the first weeks of infection. Furthermore, CRISPR XIAP KO does not significantly alter the growth or survival of established LCLs ^24^ and did not score in a human genome-wide CRISPR screen for LCL dependency factors ^40^. Therefore, the question of why EBV+ lymphomas are not observed in XLP2 patients has remained open.

We now suggest that XIAP plays key roles in support of the earliest stages of EBV-mediated primary human B-cell transformation, particularly in the presence of elevated IFNγ and TNFα, which are often found at elevated levels in XLP2 patients, who exhibit high rates of recurrent splenomegaly and HLH ^38^. Over the first three days post-infection, EBV remodels newly infected B-cells, driving elevated levels of MYC expression, altering metabolism and quadrupling their size. EBV then drives Burkitt-like B-cell hyperproliferation between days 3-7 post-infection^30,31,41–43^. This aligns with our findings, where we observed extensive cell death in EBV-infected *XIAP* knockout cells around day 4 post-infection. EBV then expresses increasing levels of the oncogene LMP1, which mimics aspects of CD40 signaling, to convert B-cells to lymphoblastoid B-cell physiology ^44^. Key LMP1/NF-kB pathway targets include anti-apoptosis factors including cIAP1, cIAP2, cFLIP and pro-survival BCL-2 family members ^45,46^. LMP1 can also inhibit p53-mediated apoptosis by inducing the deubiquitinase A20 ^47^. However, EBV does not significantly upregulate XIAP in transforming B-cells and p53 expression is sustained ^30,33^. Therefore, our results suggest that XIAP exerts key pro-survival roles likely in the Burkitt-like hyperproliferation stage of EBV transformation, prior to the marked upregulation of anti-apoptotic factors by LMP1 and perhaps also prior to maximal inhibition of p53 signaling by EBNA3C ^48,49^. Likewise, following LMP1 upregulation, lymphoblastoid cells are not dependent on XIAP for growth or survival.

What then triggers apoptosis signaling within the first week of EBV infection? EBV-driven B-cell hyper-proliferation activates DNA damage responses (DDR), which signal through p53 ^31,43,50,51^. Following the initial phase of rapid proliferation, by approximately day 7 post-EBV infection, the proliferation rate slows. Our results suggest that such elevated p53 levels and upregulation of p53 targets including NOXA and PUMA culminate in activation of BAX, which is a three BH3-only (BCL-2 Homology domain) apoptosis effector. When expressed at elevated levels and not sufficiently counteracted by pro-survival BCL2 family members, BAX undergoes conformational changes, oligomerization and insertion into the mitochondrial outer membrane. This enables egress of cytochrome c and other apoptogenic factors to activate executioner caspase activity, including caspases 3 and 7 ^34,35^. Our observations that the inhibition of p53 and BAX can avert cell death in EBV-infected XIAP-deficient cells suggest that this apoptotic pathway constitutes a pivotal mechanism of cell death during the initial stages of EBV infection. Therefore, EBV driven B-cell hyperproliferation and DNA damage signaling creates a dependency on XIAP prior to LMP1 upregulation, and this may serve to protect XLP2 B-cells from undergoing full EBV immortalization, particularly in a hyper-inflammatory cytokine milieu. This may also explain why we did not observe defects in XLP2 B-cell proliferation when stimulated by CD40L/IL-21, which highly upregulates anti-apoptotic factors including cIAP1 and 2.

How does XIAP loss of function synergize with IFNγ and TNFα to antagonize EBV-mediated B-cell transformation? P53 and IFNγ have an intricate cross-relationship, and it is plausible that IFNγ treatment heightens sensitivity to p53-driven apoptosis signaling ^52^ in newly EBV-infected B-cells. For instance, IFNγ can upregulate nuclear p53 expression and interaction with target genes ^53^. IFNγ can also block EBV-mediated B-cell transformation through incompletely identified mechanisms ^54^. TNFα may directly trigger apoptosis signaling in newly EBV-infected B-cells that may necessitate XIAP activity for survival, as is observed in XIAP KO murine bone marrow derived macrophages and in XIAP KO mice *in vivo* ^39^. We note that EBV+ lymphoblastoid cells are also dependent on cFLIP to block TNFα-mediated apoptosis ^40^. IFNγ can also induce apoptosis, and this may be exacerbated by XIAP deficiency in infected B-cells undergoing Burkitt-like hyperproliferation. Alternatively, IFNγ or TNFα may alter EBV oncogene expression to interrupt EBV-driven B-cell immortalization over the first week of infection. Identifying precise mechanisms by which IFNγ and TNFα each antagonize EBV-mediated B-cell immortalization in the absence of XIAP will be an important objective.

A recent study identified that LCLs derived from XLP2 patients had modestly elevated lytic gene expression ^24^. However, we did not observe significantly elevated lytic gene expression in our transcriptomic or proteomic profiling, suggesting that this phenomenon may be specific to established LCLs and did not likely contribute to inhibition of EBV-infected B-cell outgrowth. Similarly, the tumor suppressor cell adhesion molecule 1 (CADM1) was highly upregulated on LCLs established from XLP2 B-cells. Consistent with this report, our proteomic profiling identified CADM1 upregulation in XLP2 B-cells at 7 days post-infection (Table S2), suggesting that this is an early phenomenon in EBV-infected XLP2 B-cells.

In summary, we identified a key role for XIAP in blockade of EBV-induced apoptosis signaling within the first week of infection. Loss of XIAP function impaired proliferation and triggered apoptosis of EBV+ B-cells, particularly at the Burkitt-like stage of hyperproliferation and prior to full LMP1 upregulation and pro-survival signaling. This pro-apoptotic pathway was dependent on p53- and BAX and was exacerbated by treatment IFNγ or TNFα, whose expression are elevated with XLP2. Our results provide insights into the curious absence of EBV-driven lymphoproliferative disease in XLP2 patients, despite heightened sensitivity to EBV.

## Acknowledgments

This work was supported by NIH R01AI164709, R01CA228700, R01DE033907, U01CA275301, P01CA269043, R21AI170751 and R21AI181873 to B.E.G. and by an American Cancer Society Postdoctoral Fellowship PF-23-1144614-01-IBCD to Y.S. We thank Sam Katz (Yale University) for helpful discussions.

## Author contributions

Y.S. performed the experiments, data analysis, wrote the first draft and edited the manuscript together with B.E.G; J.C. provided blood samples of XLP-2 patients and healthy controls; K.D. performed mass spectrometry analysis; B.E.G. supervised the study. All authors read and approved the final manuscript.

## References

1. Farrell PJ. Epstein–Barr Virus and Cancer. Annual Review of Pathology: Mechanisms of Disease. 2019;14(Volume 14, 2019):29–53.

2. Münz C. Latency and lytic replication in Epstein–Barr virus-associated oncogenesis. Nature Reviews Microbiology. 2019;17(11):691–700.

3. Cohen JI, Mocarski Es Fau - Raab-Traub N, Raab-Traub N Fau - Corey L, Corey L Fau - Nabel GJ, Nabel GJ. The need and challenges for development of an Epstein-Barr virus vaccine. (1873-2518 (Electronic)).

4. Heslop HE. How I treat EBV lymphoproliferation. Blood. 2009;114(19):4002–4008.

5. Tangye SG, Latour S. Primary immunodeficiencies reveal the molecular requirements for effective host defense against EBV infection. Blood. 2020;135(9):644–655.

6. Cohen JI. Primary Immunodeficiencies Associated with EBV Disease. In: Münz C, ed. Epstein Barr Virus Volume 1: One Herpes Virus: Many Diseases. Cham: Springer International Publishing; 2015:241–265.

7. Rickinson AB, Long HM, Palendira U, Münz C, Hislop AD. Cellular immune controls over Epstein-Barr virus infection: new lessons from the clinic and the laboratory. (1471-4981 (Electronic)).

8. Parvaneh N, Filipovich AH, Borkhardt A. Primary immunodeficiencies predisposed to Epstein-Barr virus-driven haematological diseases. British Journal of Haematology. 2013;162(5):573–586.

9. Chiu Y-F, Ponlachantra K, Sugden B. How Epstein Barr Virus Causes Lymphomas. Viruses. Vol. 16; 2024.

10. Shannon-Lowe C, Rickinson AB, Bell AI. Epstein–Barr virus-associated lymphomas. Philosophical Transactions of the Royal Society B: Biological Sciences. 2017;372(1732):20160271.

11. Taylor GS, Long HM, Brooks JM, Rickinson AB, Hislop AD. The Immunology of Epstein-Barr Virus– Induced Disease. Annual Review of Immunology. 2015;33(1):787–821.

12. Meyer L, Hines M, Zhang K, Nichols KE. X-Linked Lymphoproliferative Disease. BTI - GeneReviews(®).

13. Schmid JP, Canioni D, Moshous D, et al. Clinical similarities and differences of patients with X-linked lymphoproliferative syndrome type 1 (XLP-1/SAP deficiency) versus type 2 (XLP-2/XIAP deficiency). Blood. 2011;117(5):1522–1529.

14. Henter J-I, Horne A, Aricó M, et al. HLH-2004: Diagnostic and therapeutic guidelines for hemophagocytic lymphohistiocytosis. Pediatric Blood & Cancer. 2007;48(2):124–131.

15. Coffey AJ, Brooksbank RA, Brandau O, et al. Host response to EBV infection in X-linked lymphoproliferative disease results from mutations in an SH2-domain encoding gene. Nature Genetics. 1998;20(2):129–135.

16. Sayos J, Wu C, Morra M, et al. The X-linked lymphoproliferative-disease gene product SAP regulates signals induced through the co-receptor SLAM. Nature. 1998;395(6701):462–469.

17. Hanifeh M, Ataei F. XIAP as a multifaceted molecule in Cellular Signaling. Apoptosis. 2022;27(7):441–453.

18. Rigaud S, Fondanèche M-C, Lambert N, et al. XIAP deficiency in humans causes an X-linked lymphoproliferative syndrome. Nature. 2006;444(7115):110–114.

19. Huang Y, Park YC, Rich RL, Segal D, Myszka DG, Wu H. Structural Basis of Caspase Inhibition by XIAP: Differential Roles of the Linker versus the BIR Domain. Cell. 2001;104(5):781–790.

20. Galbán S, Duckett CS. XIAP as a ubiquitin ligase in cellular signaling. Cell Death & Differentiation. 2010;17(1):54–60.

21. Filipovich AH, Zhang K, Snow AL, Marsh RA. X-linked lymphoproliferative syndromes: brothers or distant cousins? Blood. 2010;116(18):3398–3408.

22. Tangye SG. XLP: Clinical Features and Molecular Etiology due to Mutations in SH2D1A Encoding SAP. Journal of Clinical Immunology. 2014;34(7):772–779.

23. Latour S, Aguilar C. XIAP deficiency syndrome in humans. Seminars in Cell & Developmental Biology. 2015;39:115–123.

24. Engelmann C, Schuhmachers P, Zdimerova H, et al. Epstein Barr virus-mediated transformation of B cells from XIAP-deficient patients leads to increased expression of the tumor suppressor CADM1. Cell Death & Disease. 2022;13(10):892.

25. Hunte R, Alonso P, Thomas R, et al. CADM1 is essential for KSHV-encoded vGPCR-and vFLIP-mediated chronic NF-κB activation. PLOS Pathogens. 2018;14(4):e1006968.

26. Good KL, Bryant VL, Tangye SG. Kinetics of Human B Cell Behavior and Amplification of Proliferative Responses following Stimulation with IL-211. The Journal of Immunology. 2006;177(8):5236–5247.

27. Nikolovska-Coleska Z, Xu L, Hu Z, et al. Discovery of Embelin as a Cell-Permeable, Small-Molecular Weight Inhibitor of XIAP through Structure-Based Computational Screening of a Traditional Herbal Medicine Three-Dimensional Structure Database. Journal of Medicinal Chemistry. 2004;47(10):2430–2440.

28. Chou J, Platt CD, Habiballah S, et al. Mechanisms underlying genetic susceptibility to multisystem inflammatory syndrome in children (MIS-C). Journal of Allergy and Clinical Immunology. 2021;148(3):732–738.e731.

29. Philipp J, Vucic D. Regulation of Cell Death and Immunity by XIAP. Cold Spring Harbor perspectives in biology. 2020;12(8):a036426.

30. Wang LW, Shen H, Nobre L, et al. Epstein-Barr-Virus-Induced One-Carbon Metabolism Drives B Cell Transformation. Cell Metabolism. 2019;30(3):539–555.e511.

31. Nikitin PA, Yan CM, Forte E, et al. An ATM/Chk2-Mediated DNA Damage-Responsive Signaling Pathway Suppresses Epstein-Barr Virus Transformation of Primary Human B Cells. Cell Host & Microbe. 2010;8(6):510–522.

32. Pich D, Mrozek-Gorska P, Bouvet M, et al. First Days in the Life of Naive Human B Lymphocytes Infected with Epstein-Barr Virus. mBio. 2019;10(5):10.1128/mbio.01723-01719.

33. Wang C, Li D, Zhang L, et al. RNA Sequencing Analyses of Gene Expression during Epstein-Barr Virus Infection of Primary B Lymphocytes. Journal of Virology. 2019;93(13):10.1128/jvi.00226-00219.

34. Lomonosova E, Chinnadurai G. BH3-only proteins in apoptosis and beyond: an overview. Oncogene. 2008;27(1):S2–S19.

35. Aubrey BJ, Kelly GL, Janic A, Herold MJ, Strasser A. How does p53 induce apoptosis and how does this relate to p53-mediated tumour suppression? Cell Death & Differentiation. 2018;25(1):104–113.

36. Garner TP, Amgalan D, Reyna DE, Li S, Kitsis RN, Gavathiotis E. Small-molecule allosteric inhibitors of BAX. Nature Chemical Biology. 2019;15(4):322–330.

37. Komarov PG, Komarova EA, Kondratov RV, et al. A Chemical Inhibitor of p53 That Protects Mice from the Side Effects of Cancer Therapy. Science. 1999;285(5434):1733–1737.

38. Wada T, Kanegane H, Ohta K, et al. Sustained elevation of serum interleukin-18 and its association with hemophagocytic lymphohistiocytosis in XIAP deficiency. Cytokine. 2014;65(1):74–78.

39. Witt A, Goncharov T, Lee YM, et al. XIAP deletion sensitizes mice to TNF-induced and RIP1-mediated death. Cell Death Dis. 2023;14(4):262.

40. Ma Y, Walsh MJ, Bernhardt K, et al. CRISPR/Cas9 Screens Reveal Epstein-Barr Virus-Transformed B Cell Host Dependency Factors. Cell Host Microbe. 2017;21(5):580–591 e587.

41. Wang LW, Wang Z, Ersing I, et al. Epstein-Barr virus subverts mevalonate and fatty acid pathways to promote infected B-cell proliferation and survival. PLoS Pathog. 2019;15(9):e1008030.

42. Mrozek-Gorska P, Buschle A, Pich D, et al. Epstein-Barr virus reprograms human B lymphocytes immediately in the prelatent phase of infection. Proc Natl Acad Sci U S A. 2019;116(32):16046–16055.

43. McFadden K, Hafez AY, Kishton R, et al. Metabolic stress is a barrier to Epstein-Barr virus-mediated B-cell immortalization. Proc Natl Acad Sci U S A. 2016;113(6):E782–790.

44. Price AM, Tourigny JP, Forte E, Salinas RE, Dave SS, Luftig MA. Analysis of Epstein-Barr virus-regulated host gene expression changes through primary B-cell outgrowth reveals delayed kinetics of latent membrane protein 1-mediated NF-kappaB activation. J Virol. 2012;86(20):11096–11106.

45. Price AM, Dai J, Bazot Q, et al. Epstein-Barr virus ensures B cell survival by uniquely modulating apoptosis at early and late times after infection. Elife. 2017;6.

46. Mitra B, Beri NR, Guo R, Burton EM, Murray-Nerger LA, Gewurz BE. Characterization of target gene regulation by the two Epstein-Barr virus oncogene LMP1 domains essential for B-cell transformation. mBio. 2023;14(6):e0233823.

47. Husaini R, Ahmad M, Soo-Beng Khoo A. Epstein-Barr virus Latent Membrane Protein LMP1 reduces p53 protein levels independent of the PI3K-Akt pathway. BMC Res Notes. 2011;4:551.

48. Saha A, Murakami M, Kumar P, Bajaj B, Sims K, Robertson ES. Epstein-Barr virus nuclear antigen 3C augments Mdm2-mediated p53 ubiquitination and degradation by deubiquitinating Mdm2. J Virol. 2009;83(9):4652–4669.

49. Yi F, Saha A, Murakami M, et al. Epstein-Barr virus nuclear antigen 3C targets p53 and modulates its transcriptional and apoptotic activities. Virology. 2009;388(2):236–247.

50. Nikitin PA, Price AM, McFadden K, Yan CM, Luftig MA. Mitogen-induced B-cell proliferation activates Chk2-dependent G1/S cell cycle arrest. PLoS One. 2014;9(1):e87299.

51. Koganti S, Hui-Yuen J, McAllister S, et al. STAT3 interrupts ATR-Chk1 signaling to allow oncovirus-mediated cell proliferation. Proc Natl Acad Sci U S A. 2014;111(13):4946–4951.

52. Ossina NK, Cannas A, Powers VC, et al. Interferon-gamma modulates a p53-independent apoptotic pathway and apoptosis-related gene expression. J Biol Chem. 1997;272(26):16351–16357.

53. Contreras AU, Mebratu Y, Delgado M, et al. Deacetylation of p53 induces autophagy by suppressing Bmf expression. J Cell Biol. 2013;201(3):427–437.

54. Lunemann A, Vanoaica LD, Azzi T, Nadal D, Munz C. A distinct subpopulation of human NK cells restricts B cell transformation by EBV. J Immunol. 2013;191(10):4989–4995.

