## Supplemental materials for "Insights into the Absence of Lymphoma Despite Fulminant Epstein-Barr Virus Infection in Patients with XIAP Deficiency"

### Supplementary Figure legends and tables

**Figure S1. CRISPR XIAP KO does not alter the growth or survival of GM15829 LCLs.**

(A) FACS analysis of primary human B cells electroporation efficiency. Shown are FACS plots of primary human B-cells at 1-hour post-electroporation with control unlabeled versus ATTO™ 550 conjugated tracrRNA duplex.

(B) Immunoblot analysis of WCL from Cas9+ GM15892 LCLs that expressed the indicated control or one of four independent XIAP-targeting sgRNAs.

(C) Growth curve analysis of Cas9+ GM15892 cells expressing control or *XIAP* targeting sgRNAs. Shown are mean ± SD fold change live cell numbers from n=3 replicates, relative to Day 0 values.

Statistical significance was assessed by one-way ANOVA followed by Tukey's multiple comparisons test (C). Blots are representative of n=3 replicates. ns, not significant.

**Figure S2. Outgrowth of XLP2 cells is impaired at early timepoints of EBV infection but not of CD40L/IL-21 treatment.**

(A) Schematic diagram highlighting the XIAP mutation shared by XLP2 Patients #3 and #4.

(B) Growth curve analysis of primary B-cells from XLP2 patients versus controls following EBV infection. Shown are mean ± SD fold change live cell numbers from n=3 replicates of B-cells infected at Day 0 with EBV, relative to Day 0 values. The annotations represent the results of statistical comparisons between XLP2 samples and Control #4.

(C) Growth curve analysis of primary B-cells from XLP2 patients versus controls stimulated by CD40L/IL-21. Shown are mean ± SD fold change live cell numbers from n=3 replicates of B-cells stimulated by CD40L and IL-21, which were replenished every 3 days. The annotations represent the results of statistical comparisons between XLP2 samples and Control #4.

Statistical significance was assessed by one-way ANOVA followed by Tukey's multiple comparisons test (B and C). *, $p<0.05$; **, $p<0.01$; ***, $p<0.001$; ****, $p<0.0001$; ns, not significant.

**Table S1. sgRNA sequence used in this study**

| sgRNA name | sgRNA sequence |
|---|---|
| Control | ATTTCGCAGATCATCGACAT |
| XIAP #1 | GCATCAACACTGGCACGAGC |
| XIAP #2 | AGTGCTGGACTCTACTACAC |
| XIAP #1 | ATGACAACTAAAGCACCGCA |
| XIAP #2 | ATGGATATACTCAGTTAACA |

| | |
|---|---|
| XIAP #3 | TCTGACCAGGCACGATCACA |
| XIAP #4 | TATCAGACACCATATACCCG |

**Supplementary Material & Methods**

Cell lines

HEK293T cells were cultured in DMEM medium containing 10% fetal bovine serum (FBS). GM12878 and GM15892 Cas9[+] lymphoblastoid cell lines were cultured in RPMI-1640 supplemented with 10% FBS and 5 µg/ml blasticidin. All cells were incubated in a humidified incubator at 37°C with 5% CO2 and were routinely confirmed to be mycoplasma-negative using the MycoAlert kit (Lonza).

Chemical compounds

Proliferation of primary B-cells was induced by 50 ng/mL rhCD40L (Enzo Life Sciences) plus 50 ng/mL rhIL-21 (Biolegend). For the inhibition of XIAP, 5 µM Embelin (Selleck) was used. 20 µM zVAD-Fmk (Selleck) was used to inhibit caspase 3 and 7. 5 µM BAI1 (MedChemexpress) and 5 or 20 µM Pifithrin-α (MedChemexpress) were used for inhibition of Bax and p53, respectively. All reagents were replenished every 3 days.

Immunoblot analysis

Cells were lysed in 1× Laemmli buffer with ultrasound, and then boiled at 95°C for 10 minutes. The proteins were separated by SDS-PAGE electrophoresis and transferred onto the nitrocellulose membranes. The membrane was blocked with 5% milk in TBST buffer and incubated with primary antibodies at 4°C overnight, followed by 1 hour incubation with secondary antibodies at room temperature.  Blots were then developed by incubation with ECL chemiluminescence for 1 min (Millipore) and images were captured by Licor Fc platform.

Flow cytometry analysis

For live cells staining, 1× $10^6$ of cells were washed twice with FACS buffer (PBS containing 1% FBS), followed by primary antibody incubation at dark for 30 minutes. For 7-AAD staining, cells were stained with 7-AAD for 10 minutes on ice. Then the stained cells were washed with FACS buffer and subjected to flow cytometry analysis. Data was analyzed with FlowJo X software (FlowJo).

Proliferation assay

Cells were seeded in a 12 well plate at 0.5 million cells per well and treated with indicated reagents. Culture medium was changed every 3 or 4 days, and the live cell numbers were counted on indicated days using trypan blue staining. Cell counts relative to day 0 were calculated and used for the plots.

CFSE and Cell trace violet staining

For CFSE or cell trace violet staining, cells were washed twice with PBS, followed by staining with 1 μM CFSE or cell trace violet at dark for 20 mins. Then the cells were washed twice with RPMI-1640 containing 10% FBS and subjected to subsequent experiments.

Caspase-Glo 3/7 Assay

Caspase 3/7 activity was assessed using Caspase-Glo® 3/7 Assay (Promega) according to manufacturer’s instructions. In brief, cells were mixed with Caspase-Glo® 3/7 reagent at a ratio of 1:1 and incubated for 1 hour at room temperature. The luminescence signal was assessed with a Molecular Devices microplate reader. The final presentation of caspase-3/7 activities was established by normalizing the signal against the count of viable cells, determined through trypan blue staining.

RNAseq analysis

Total RNA was extracted from the cells using RNeasy Mini kit (Qiagen), following the manufacturer's instruction. A DNA digestion step within the column was incorporated to eliminate any remaining genomic DNA contamination. For library construction, poly(A) mRNA was isolated from 1 µg DNA-free total RNA using NEBNext Poly(A) mRNA Magnetic Isolation Module (New England Biolabs), followed by library construction via the NEBNext Ultra RNA Library Prep Kit (New England Biolabs). Library quality was assessed by an Agilent Bioanalyzer DNA Chip. Multi-indexed libraries were combined, pooled, and subjected to sequencing on an Illumina NextSeq 500 sequencer with single-end 75 bp reads at the Dana Farber Molecular Biology core. Raw read counts for gene expression were quantified using salmon v1.2.0 with human GENCODE v28 (GRCh37) genes. Differential expressions were evaluated by DESeq2.

Mass spectrometry analysis

Cells were lysed in (50 mM Tris-HCl pH 7.5, 300 mM NaCl, 0.5% v/v NP40, 1 mM DTT and Roche protease inhibitor cocktail). Proteins were precipitated with 20% trichloroacetic acid (TCA), washed once with 10% TCA, washed three times with cold acetone and dried to completion, using a centrifugal evaporator. Samples were resuspended in digestion buffer (50 mM Tris-HCl pH 8.5, 10% acetonitrile (AcN), 1 Mm DTT, 10 mg/ml trypsin (Promega) and incubated overnight at 37oC, with agitation. The reaction was quenched by 50% formic acid (FA), subjected to C18 solid-phase extraction, and vacuum-centrifuged to complete dryness. Samples were reconstituted in 4% AcN/5% FA and divided into technical duplicates prior to LC-MS/MS on an Orbitrap Lumos.

Data availability

Requests for resources and reagents can be directed to the first author or corresponding author. Figures were drawn with commercially available GraphPad, Biorender and Microsoft Powerpoint.

### Statistics

Unless otherwise indicated, all bar graphs and line graphs represent the arithmetic mean of three independent experiments ($n = 3$), with error bars denoting standard deviations. The statistical significance between different groups was assessed using the unpaired Student's t-test or analysis of variance (ANOVA) with the appropriate post-test using GraphPad Prism 9 software. P values correlate with symbols as follows, ns = not significant, $p > 0.05$; *$p < 0.05$; **$p < 0.01$; ***$p < 0.001$, ****$p < 0.0001$.

**Figure S1**

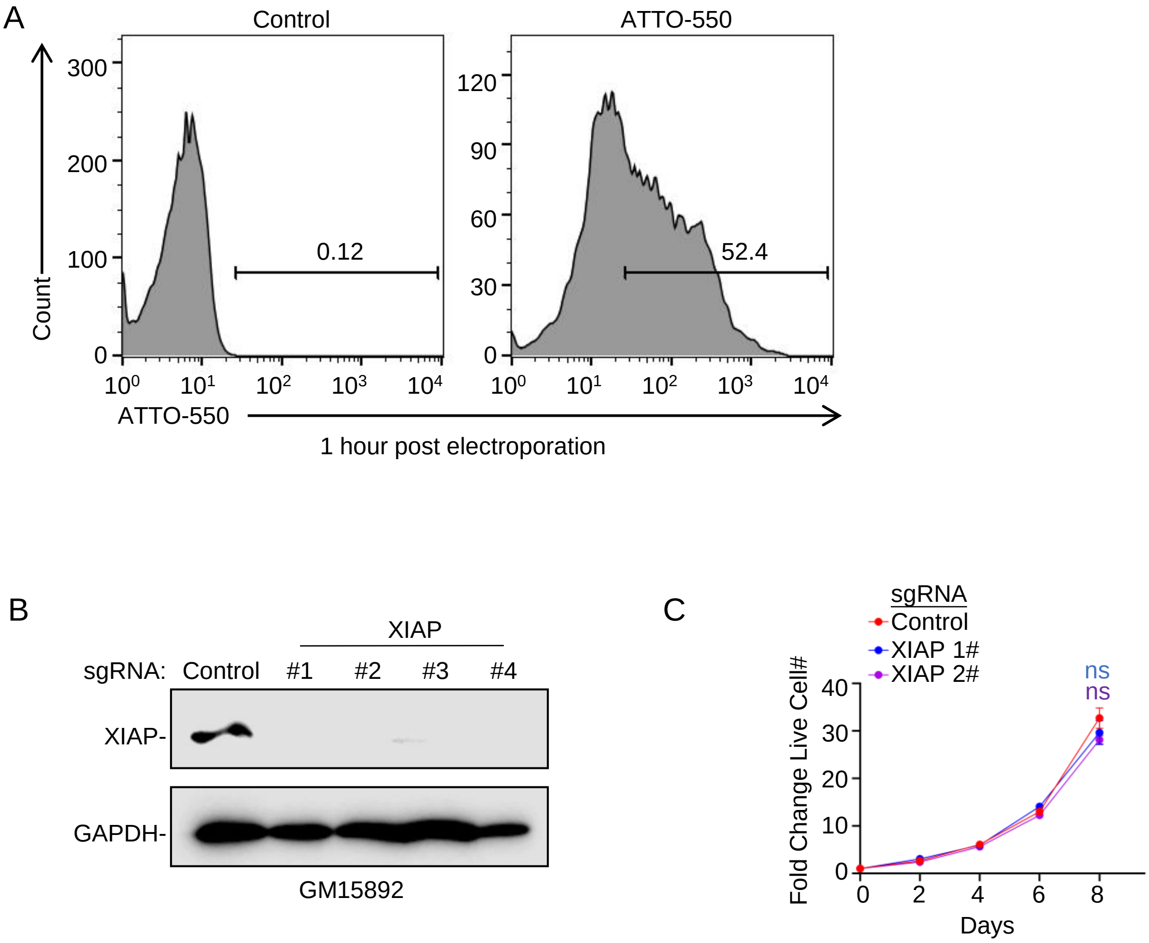
A
Control
ATTO-550
Count
300
200
100
0
120
90
60
30
0.12
52.4
$10^0$ $10^1$ $10^2$ $10^3$ $10^4$
ATTO-550
1 hour post electroporation
B
XIAP
sgRNA: Control #1 #2 #3 #4
XIAP-
GAPDH-
GM15892
C
sgRNA
Control
XIAP 1#
XIAP 2#
ns
ns
Fold Change Live Cell#
40
30
20
10
0
0 2 4 6 8
Days

Figure S1. CRISPR XIAP KO does not alter the growth or survival of GM15829 LCLs.

(A) FACS analysis of primary human B cells electroporation efficiency. Shown are FACS plots of primary human B-cells at 1-hour post-electroporation with control unlabeled versus ATTO™ 550 conjugated tracrRNA duplex.

(B) Immunoblot analysis of WCL from Cas9+ GM15892 LCLs that expressed the indicated control or one of four independent XIAP-targeting sgRNAs.

(C) Growth curve analysis of Cas9+ GM15892 cells expressing control or XIAP targeting sgRNAs. Shown are mean ± SD fold change live cell numbers from n=3 replicates, relative to Day 0 values.

Statistical significance was assessed by one-way ANOVA followed by Tukey's multiple comparisons test (C). Blots are representative of n=3 replicates. ns, not significant.

**Figure S2**

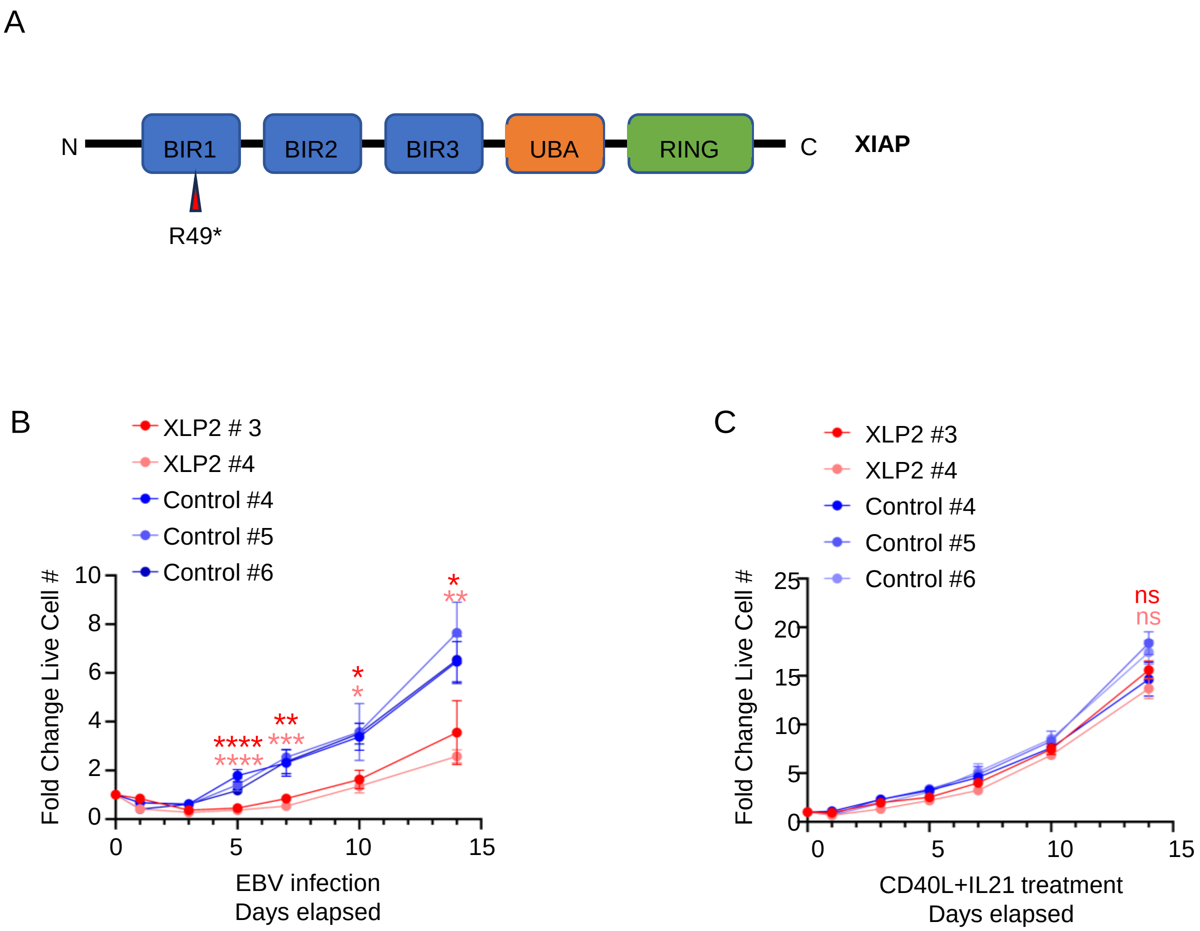
A
N
BIR1
BIR2
BIR3
UBA
RING
C
XIAP
R49*
B
XLP2 # 3
XLP2 #4
Control #4
Control #5
Control #6
Fold Change Live Cell #
EBV infection
Days elapsed
C
XLP2 #3
XLP2 #4
Control #4
Control #5
Control #6
ns
ns
Fold Change Live Cell #
CD40L+IL21 treatment
Days elapsed

Figure S2. Outgrowth of XLP2 cells is impaired at early timepoints of EBV infection but not of CD40L/IL-21 treatment.

(A) Schematic diagram highlighting the XIAP mutation shared by XLP2 Patients #3 and #4.

(B) Growth curve analysis of primary B-cells from XLP2 patients versus controls following EBV infection. Shown are mean ± SD fold change live cell numbers from n=3 replicates of B-cells infected at Day 0 with EBV, relative to Day 0 values. The annotations represent the results of statistical comparisons between XLP2 samples and Control #4.

(C) Growth curve analysis of primary B-cells from XLP2 patients versus controls stimulated by CD40L/IL-21. Shown are mean ± SD fold change live cell numbers from n=3 replicates of B-cells stimulated by CD40L and IL-21, which were replenished every 3 days. The annotations represent the results of statistical comparisons between XLP2 samples and Control #4.

Statistical significance was assessed by one-way ANOVA followed by Tukey's multiple comparisons test (B and C). *, $p<0.05$; **, $p<0.01$; ***, $p<0.001$; ****, $p<0.0001$; ns, not significant.
